# Taste adaptations associated with host-specialization in the specialist *Drosophila sechellia*

**DOI:** 10.1101/2022.11.21.517453

**Authors:** Carolina E. Reisenman, Joshua Wong, Namrata Vedagarbha, Catherine Livelo, Kristin Scott

**Affiliations:** Department of Molecular and Cell Biology, University of California Berkeley, USA; Department of Integrative Biology, University of California Berkeley, USA; Essig Museum of Entomology, University of California, USA; Design Therapeutics, Carlsbad, California, USA

**Keywords:** *Drosophila*, specialization, taste, gustatory receptors, behavior, Noni, *Drosophila sechellia*

## Abstract

Chemosensory-driven hostplant specialization is a major force mediating insect ecological adaptation and speciation. *Drosophila sechellia*, a species endemic to the Seychelles islands, feeds and oviposits on *Morinda citrifolia* almost exclusively. This fruit is harmless to *D. sechellia* but toxic to other *Drosophilidae*, including the closely related generalists *D. simulans* and *D. melanogaster*, due to its high content of fatty acids. While several olfactory adaptations mediating *D. sechellia’s* preference for its host have been uncovered, the role of taste has been much less examined. We found that *D. sechellia* has reduced taste and feeding aversion to bitter compounds and host fatty acids that are aversive to *D. melanogaster* and *D. simulans*. The loss of aversion to canavanine, coumarin, and fatty acids arose in the *D. sechellia* lineage, as its sister species *D. simulans* showed responses akin to those of *D. melanogaster. D. sechellia* has increased taste and feeding responses towards *M. citrifolia*. These results are in line with *D. sechellia’s* loss of genes encoding bitter gustatory receptors (GRs) in *D. melanogaster*. We found that two *GR* genes which are lost in *D. sechellia*, *GR39a.a* and *GR28b.a*, influence the reduction of aversive responses to some bitter compounds. Also, *D. sechellia* has increased appetite for a prominent host fatty acid compound that is toxic to its relatives. Our results support the hypothesis that changes in the taste system, specifically a reduction of sensitivity to bitter compounds that deter generalist ancestors, contribute to the specialization of *D. sechellia* for its host.

**Summary statement:** Taste specializations in the specialist *Drosophila sechellia* include a lineage-specific reduced sensitivity to bitter compounds associated with losses of gustatory receptors, and increased appetite for Noni and host fatty acids.

## INTRODUCTION

Chemosensory-driven hostplant specialization is a major force mediating insect ecological adaptation and speciation (Berlocher and Feder, 2002; Chapman, 2003; Jaenike, 1990). The role of the olfactory sensory system in mediating these processes is well understood in many species (Hansson and Stensmyr, 2011; Schoonhoven et al., 2005; Zhao and McBride, 2020). For instance, plant volatiles contribute to sympatric speciation and reproductive isolation via hostplant shifts, as in the larch bud moth *Zyraphera diniana* (Emelianov et al., 2003; Syed et al., 2003) and in the apple maggot *Rhagoletis pomonella* (Linn Jr et al., 2003; Olsson et al., 2009). Whether the taste system also contributes to host specialization has been less studied, despite its essential role for recognition and acceptance of food and oviposition sources. Genomic studies suggest that in general insects with specialized diets underwent losses/pseudogenizations in genes mediating taste detection relative to generalists (Anholt, 2020; Cande et al., 2013; Robertson, 2019), but investigations on the functional consequences are relatively scarce.

The drosophilid fly *Drosophila sechellia* is endemic to the Seychelles Islands in the Indian Ocean, which feeds and oviposits on the fruit of *Morinda citrifolia* (Lachaise et al., 1988; Tsacas and Bachli, 1981), known as “Noni”. Although a Noni specialist, flies could occasionally be found in other fruits such as mango, figs and papaya (Matute and Ayroles, 2014; Salazar-Jaramillo and Wertheim, 2021). *D. sechellia* has been much studied because it is a specialist closely related to the generalist saprophagous *D. melanogaster* and *D. simulans*, providing an excellent opportunity for studying the genetic, physiological and behavioral mechanisms underlying host specialization (Auer et al., 2022; Dekker et al., 2006; Stensmyr et al., 2003; Zhao and McBride, 2020). *D. sechellia* has a common ancestor with these species about 3-5 mya and 0.25 mya respectively (Garrigan et al., 2012). While *D. sechellia* uses Noni as its host, its relatives are repelled due to the high content of hexanoic and octanoic acid (19% and 58% of respectively) in the ripe (but not green or rotten) fruit (Farine et al., 1996; Pino et al., 2010; R’Kha et al., 1991; although *D. simulans* can be found occasionally on Noni in the Seychelles, Matute and Ayroles, 2014). Specialization on Noni might also provide protection from wasp parasitism, as these fatty acids are toxic to the larvae (Salazar-Jaramillo and Wertheim, 2021). At >1% vol/vol, octanoic acid is toxic (via smell and contact) to many drosophilids, including *D. simulans* and *D. melanogaster* (Farine et al., 1996; Legal et al., 1994). In contrast, *D. sechellia* evolved detoxification mechanisms to cope with octanoic and hexanoic acid (Drum et al., 2022; Lanno et al., 2019; Legal et al., 1994; Pino et al., 2010), and these furthermore stimulate oviposition and egg production (Álvarez-Ocaña et al., 2022; Amlou et al., 1998; Jones, 2005; Lavista-Llanos et al., 2014; R’Kha et al., 1991). At <1% vol/vol, food solutions containing these fatty acids are preferred over those lacking them, but this preference is stronger in *D. sechellia* than in *D. melanogaster* (Ferreira et al., 2020).

Insects detect chemicals in the air (i.e. by olfaction) and by contact (i.e. by taste) using specialized chemosensory cells housed in cuticular structures called sensilla. Gustatory sensilla are found mostly in the proboscis and the leg tarsi (Chapman, 1998; Chen and Dahanukar, 2020; Scott, 2018; Vosshall and Stocker, 2007). These sensilla house gustatory receptor neurons (GRNs) responsive to sweet, low salt, high salt, bitter, sour and water compounds (Chen and Dahanukar, 2020; French et al., 2015a; Liman et al., 2014; Scott, 2018). Caloric substances activate sweet GRNs triggering proboscis extension and ingestion, while bitter (potentially toxic) substances produce aversion by activating bitter GRNs; some bitter substances additionally suppress sugar GRN activation (French et al., 2015b; Jeong et al., 2013). GRNs express several chemosensory proteins, including gustatory receptors (GRs; Fujii et al., 2015; Robertson et al., 2003; Scott et al., 2001; Thorne et al., 2004), ionotropic receptors (IRs, Benton et al., 2009; Croset et al., 2010; Koh et al., 2014; Ni, 2021), and odorant binding proteins (OBPs, Galindo and Smith, 2001). GRNs express multiple chemosensory proteins (Chen and Dahanukar, 2020; Montell, 2021).

Some of the chemosensory specializations of *D. sechellia* for its host involve changes in the olfactory system that increase long-distance attraction to Noni volatiles, such as increases in the number of olfactory sensilla for detection of the signature host-compounds ethyl and methyl hexanoate, and molecular changes in OR22a and IR75b that mediate olfactory attraction to them (Auer et al., 2020, 2021; Dekker et al., 2006; Prieto-Godino et al., 2017; Stensmyr, 2009; Zhao and McBride, 2020). Hostplant taste specializations have been less explored. Matsuo et al. (2007) found that losses of OBP57d and OBP57e mediate contact-mediated oviposition acceptance of fatty acids, although both olfactory and gustatory inputs are required for oviposition on Noni substrates (Álvarez-Ocaña et al., 2022). Genomic studies showed that *D. sechellia* lost many *GR* and “divergent” *IR* genes typically expressed in external GRNs (Crava et al., 2016; Croset et al., 2010; McBride, 2007; McBride and Arguello, 2007) which detect bitter compounds and possibly fatty acids in the generalist *D. melanogaster* (Ahn et al., 2017; Brown et al., 2021; Dweck and Carlson, 2019; Masek and Keene, 2013; Sánchez-Alcañiz et al., 2018; Scott, 2018). Mc Bride and Arguello (2007) suggested two explanations for the losses of chemosensory genes: 1) after specialization on *M. citrifolia*, *D. sechellia* was exposed to fewer bitter (likely harmful) compounds in their ecological niche, leading to loss of selection for chemosensory genes; 2) the loss of chemoreceptors for detection of *M. citrifolia* deterrents facilitated the specialization of *D. sechellia* on *M. citrifolia*. In line with these predictions, electrophysiological studies found major losses of sensitivity to some bitter compounds (Dweck and Carlson, 2020), and *D. sechellia* has increased feeding preference for solutions containing host fatty acids, in comparison with its close relatives (Ferreira et al., 2020). Thus, these findings suggest that the taste is also involved in hostplant adaptation and specialization in *D. sechellia*.

Here we used behavioral assays to further study the taste and feeding responses of *D. sechellia* and its generalist relatives. We found that *D. sechellia* has a reduced behavioral aversion to bitter compounds, which correlates with the lineage-specific loss of two *GR* genes (*GR39a.a* and *GR28b.A*), and increased appetite for Noni fatty acids. Our findings thus support the hypothesis that host specialization in this fly involves adaptive changes in the taste system.

## MATERIALS AND METHODS

### Animals

*D. sechellia* flies (strain 14021-0248.27, provided by the former University of California San Diego stock center, and strain 14021-0248.25, provided by Dr. M. Eisen) and *D. melanogaster* (strain Canton-S) were used in most experiments. *D. sechellia* has low genetic diversity, but it is proposed that its small effective population size resulted from trade-offs between life history traits and the use of a predictable competition-free host (Legrand et al. 2009). All experiments were conducted with mated female flies (and a few with *D. sechellia* males). Two additional lines of *D. simulans* (strains 14021-0251.312 and 14021-0251.269) and *D. melanogaster* (strain 14021-0231.199) were obtained from the former UCSD stock center. Other *D. melanogaster* lines used include *D. melanogaster* null *GR28b.A* mutant (obtained from the Bloomington Stock Center; RRID:BDSC 24190), strain *w118;* and wCS:GR39a, wCS; generously provided by Dr. J. Carlson (Yale University). *D. melanogaster* and *D. simulans* flies were grown on standard fly food at room temperature. *D. sechellia* flies were reared at 25 °C on standard fly food supplemented with a small piece of *M. citrifolia* fruit leather (Hawaiian Organic Noni LLC) and a pinch of dry yeast. Flies were 2-7 days old at the time of the experiments.

### Behavioral assays

#### Proboscis extension response (PER) and temporal consumption assays (TCA)

Assays were performed on individual mated flies which were food-deprived by placing them in vials containing two pieces of water-saturated tissue paper. After 22-24 hours, flies were gently anesthetized under CO2 and glued by their dorsal thorax onto a glass slide using clear nail polish. Flies were allowed to recover in a humid chamber for 2 hours before PER/TCA tests. Mounted flies were first water-satiated and for PER assays, either tarsi from all six legs or the proboscis of individual animals was stimulated three times at five seconds intervals with a drop of an appetitive solution (1 M sucrose, Fisher Chemicals, Waltham, Massachusetts, USA, CAS # 57-50-1; or 1 M D-glucose, Sigma-Aldrich, Saint Louis, Missouri, USA, CAS # 59-99-7), a mixture of 1 M sucrose and 0.5 mM denatonium, or Noni fruit leather strips (Hawaiian organic Noni LLC, Kauai, Hawaii, USA) reconstituted by blending it with water (0.00657 g/ml), and the number of proboscis extensions was recorded (Fig. 1A). The number of animals that did or did not extend their proboscis was calculated and data was analyzed using Fisher Exact tests (for comparisons between independent groups) or McNemar tests (for paired data) (Zar, 1999). Data was considered significant if p<0.05. In subsequent experiments we used glucose as a sugar stimulus because it evokes stronger responses than sucrose in *D. sechellia* (Fig. 1B-C), and because it naturally occurs in Noni, although in small amounts (Potterat and Hamburger, 2007).

**Figure 1:**
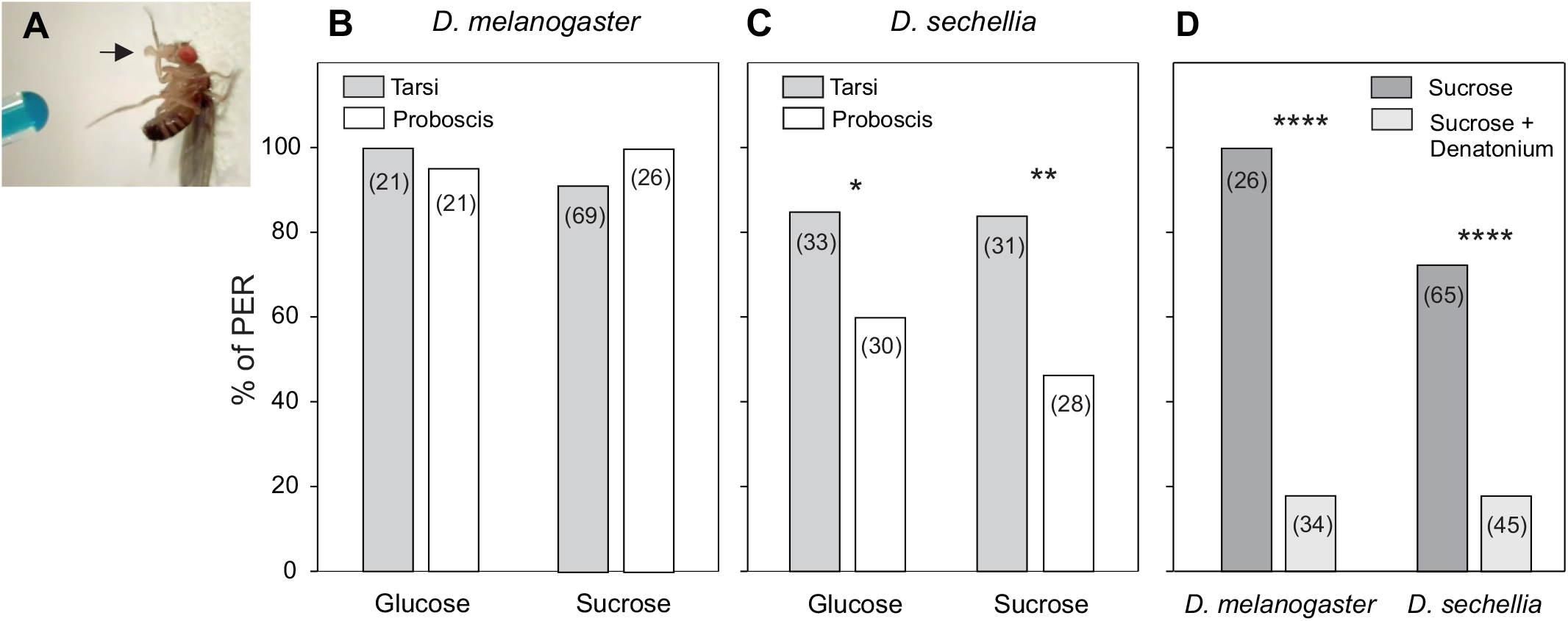
PER responses are organ- and species-specific. **(A)** Schematic of the fly preparation. Food-deprived female flies were mounted in a glass slide, and the taste organs were stimulated with food solution (here dyed blue for visualization) applied to all leg tarsi or the proboscis; flies were not allowed to drink. Each fly was tested with one condition only. Data show the proportion of flies that extend their proboscis at least once; numbers between parentheses indicate the number of flies tested. **B:** In *D. melanogaster*, PER to 1 M sugar was independent of the organ stimulated (p>0.05), but higher upon proboscis stimulation at lower concentrations; Supp. Fig. 1A). **C:** In *D. sechellia* PER was higher upon tarsi stimulation (Fisher Exact tests, *p<0.05, **p<0.01). **C:** Stimulation with 1 M sucrose + a bitter compound (0.5 mM Denatonium) reduced PER in both species (***p<0.005); data obtained from different animals than in **A-B.**

TCA were performed similarly to those reported by Yao and Scott (2022). Two-four days old mated females were food-deprived for 22 hours and prepared for experiments as above (Fig. 1A). All tarsi from individual flies were touched with a drop of 750 mM glucose or Noni juice (Noni only, Dynamic Health Laboratories, New York, New York, USA), as the fruit juice constitutes a more reproducible stimulus than the ripe fruit (Auer et al., 2020). Each fly was stimulated up to 10 consecutive times and allowed to drink, and then stimulated again in the same fashion until flies stopped consuming (usually before 60 seconds). The time and duration of each feeding event was manually recorded using an online chronometer (http://online-stopwatch.chronme.com/). Data was exported and the following parameters were calculated off-line for each fly and stimulus (Figure 2C): the number of feeding events, the total feeding time (summed duration of all feeding events), and the percentage of animals that fed.

**Figure 2:**
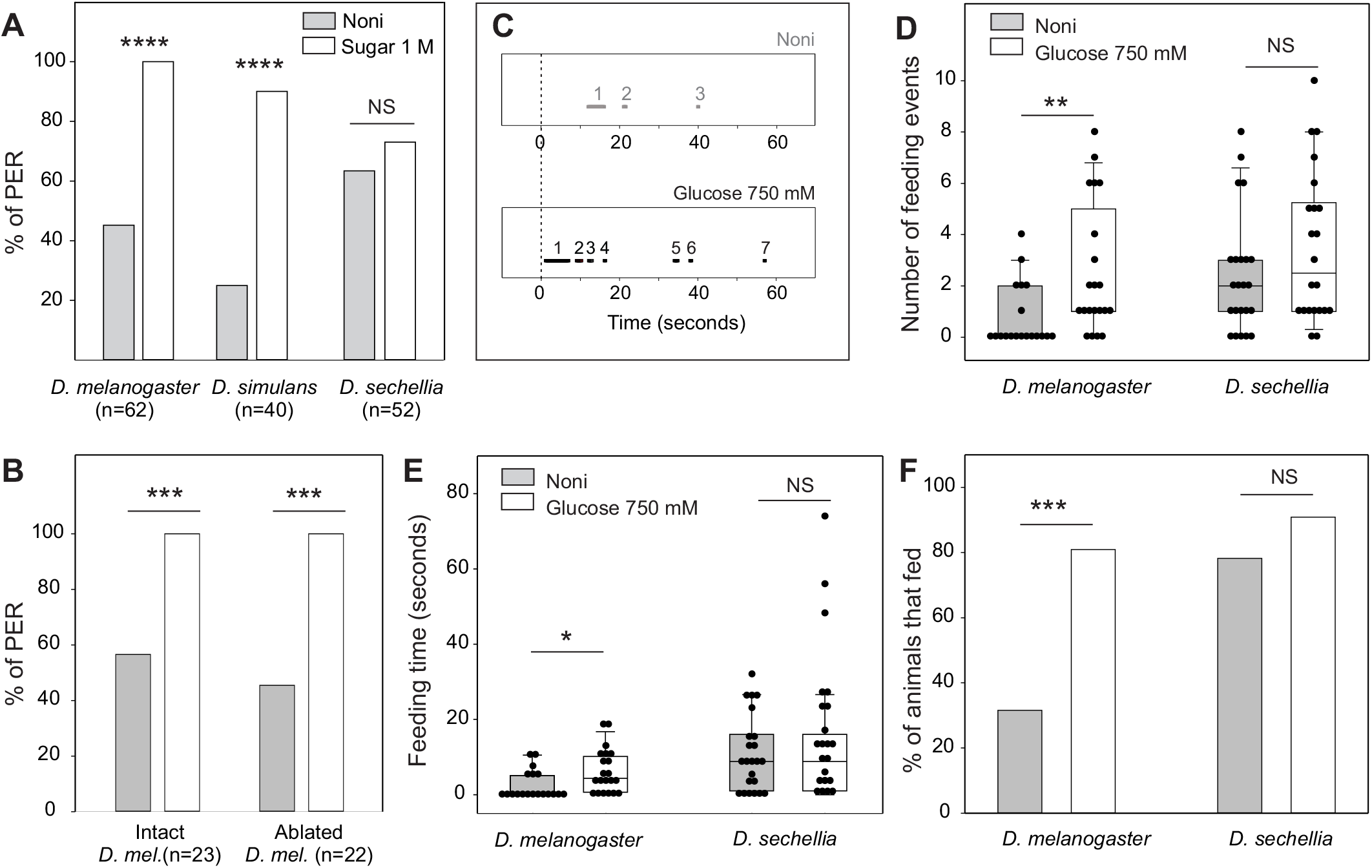
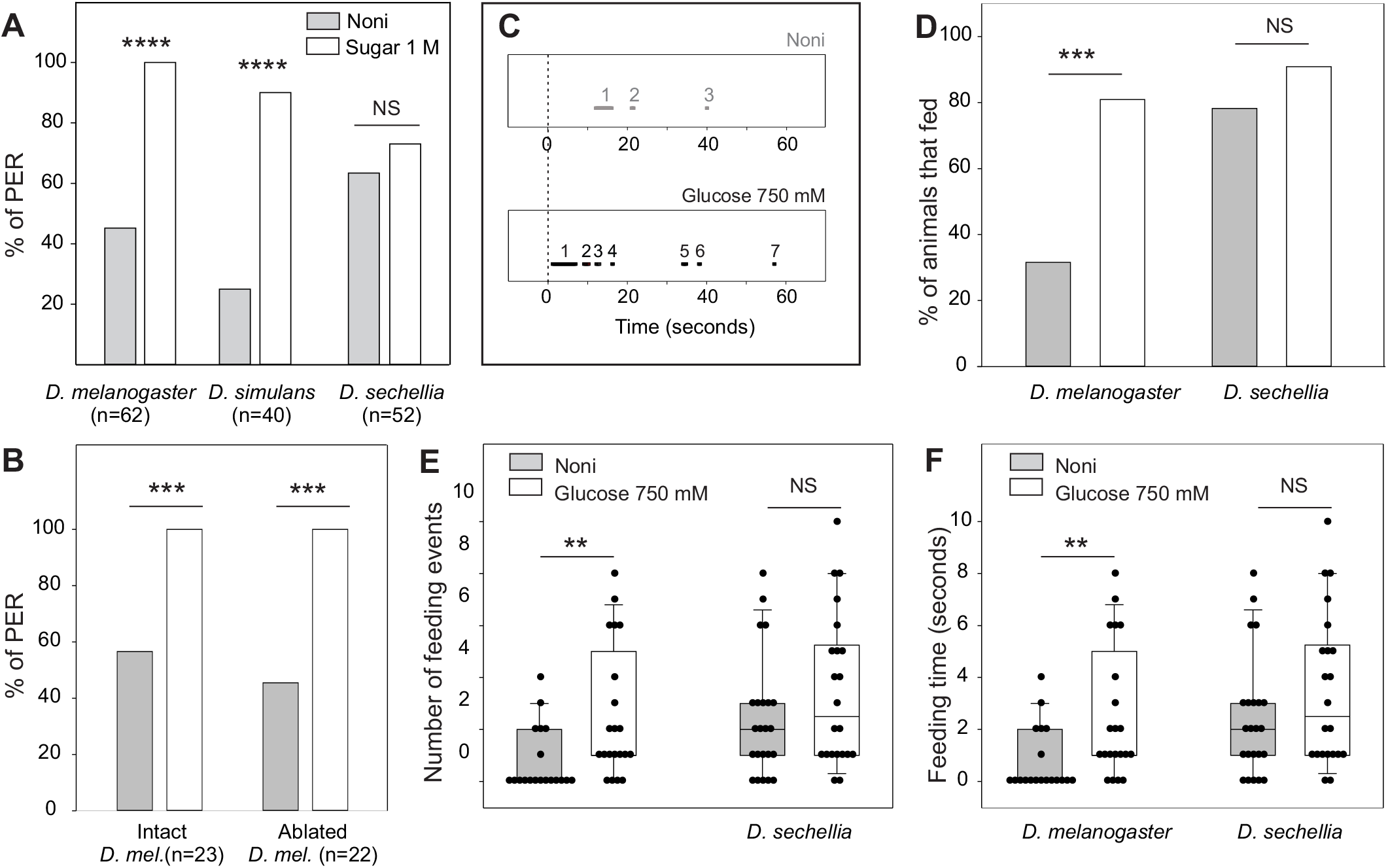
*D. sechellia* has comparatively increased taste and feeding responses to Noni. **A:** PER across species. Flies were prepared as in Fig. 1A and their tarsi were stimulated three times first with 1 M sugar solution (sucrose for the generalists and glucose for *D. sechellia* given their differential responsiveness, Figs. 1C and S1B), and then with Noni. Flies were not allowed to drink sugar or Noni, but could drink water between presentation of these food solutions. The proportion of flies showing PER to sugar stimulation was higher than that of animals showing PER to Noni in *D. melanogaster* and *D. simulans* (Mc Nemar tests, ***p<0.005 in both cases), while *D. sechellia* showed similar PER to both stimuli (NS, p>0.05). **B:** Reduced PER to Noni in the generalists does not require olfaction. Parallel cohorts of intact and anosmic flies (olfactory organs ablated two days before tests) were assayed as in **A;** both groups showed similarly reduced PER to Noni in comparison with Sucrose 1 M (***p<0.005 in both cases). **C-E:** Temporal consumption assay of 24-hours food-deprived restrained flies. Flies were prepared as in Fig. 1A; their tarsi were stimulated up to 10 consecutive times with 750 mM glucose or Noni juice and allowed to drink. We stimulated tarsi because it evoked stronger responses than proboscis stimulation (Fig. 1). Timing of feeding responses were digitally recorded and analyzed off-line. Here and in all upcoming experiments, we used glucose instead of sucrose because *D. sechellia* has a higher responsiveness to this sugar (which occurs in Noni). **C:** Examples of the temporal sequence of ingestion of individual *D. melanogaster* flies offered Noni or glucose. The dotted vertical lines at zero indicate the beginning of the tests. Black bars indicate feeding events (3 and 7 respectively); the summed duration of all feeding events was 5 and 10 sec in these examples. **D**: In *D. melanogaster*, the percentage of flies that consumed Noni was lower than that of flies that consumed glucose (asterisks, Fisher Exact test, p<0.005; n=19, 21), but not in *D. sechellia* (NS, p>0.05; n=23-22). **E-F:** The number of feeding events and the feeding duration was higher in *D. melanogaster* when glucose was offered (left, Mann-Whitney U tests, *p<0.05, **p<0.01), but were similar upon stimulation with either stimulus in *D. sechellia* (right, p>0.05, NS). Symbols are individual data, boxes indicate the 25% and 75% quartiles, the horizontal line inside boxes indicate the median, and the whiskers indicate the 10 and 90% quartiles (this data representation is used in all figures thereafter).

#### Feeding assay

Feeding assays were conducted similarly to those described previously (Reisenman and Scott, 2019). Groups of 2-7 days old mated flies (n=11-15) were food-deprived for 24 hours, and then transferred to a vial containing a piece of filter paper (2.7 cm diameter, Whatman, cat. No 1001 125) impregnated with 160 μl of food solution dyed blue with erioglaucine (0.25 mg/ml, Sigma-Aldrich, Saint Louis, Missouri, USA, CAS # 3844-45-9). To facilitate feeding, vials were flipped upside down so that the filter paper with food solution faced up (Figure 3A). Flies had access to food for 30 minutes and then vials were frozen for at least 60 minutes. After freezing, flies in each vial were individually scored, blind to treatment, using the following five-point scale based on the amount of food visualized as blue dye in the abdomen: 0 (no dye=no food), 0.25 (“trace” of blue dye =“taste” of food), 0.5 (up to ¼ of the abdomen dyed blue), 1 (more than ¼ but less than ½ of the abdomen dyed blue), and 2 (more than ½ of the abdomen dyed blue) (Figure 3A). For each vial a single feeding score value was calculated as: (0 x n_0_ + 0.25 x n_0.25_ + 0.5 x n_0.5_ + 1 x n_1_ + 2 x n_2_ / N), where n_(0-2)_ denotes the number of flies in each score category, and N the total number of flies/vial; this single feeding score constituted a biological replicate (Reisenman and Scott, 2019). All experiments and scoring were conducted blind to treatment. In one experiment single flies (i.e., a single fly/vial) had access to food for 30 minutes, were visually inspected for viability every five minutes, frozen, and scored.

**Figure 3:**
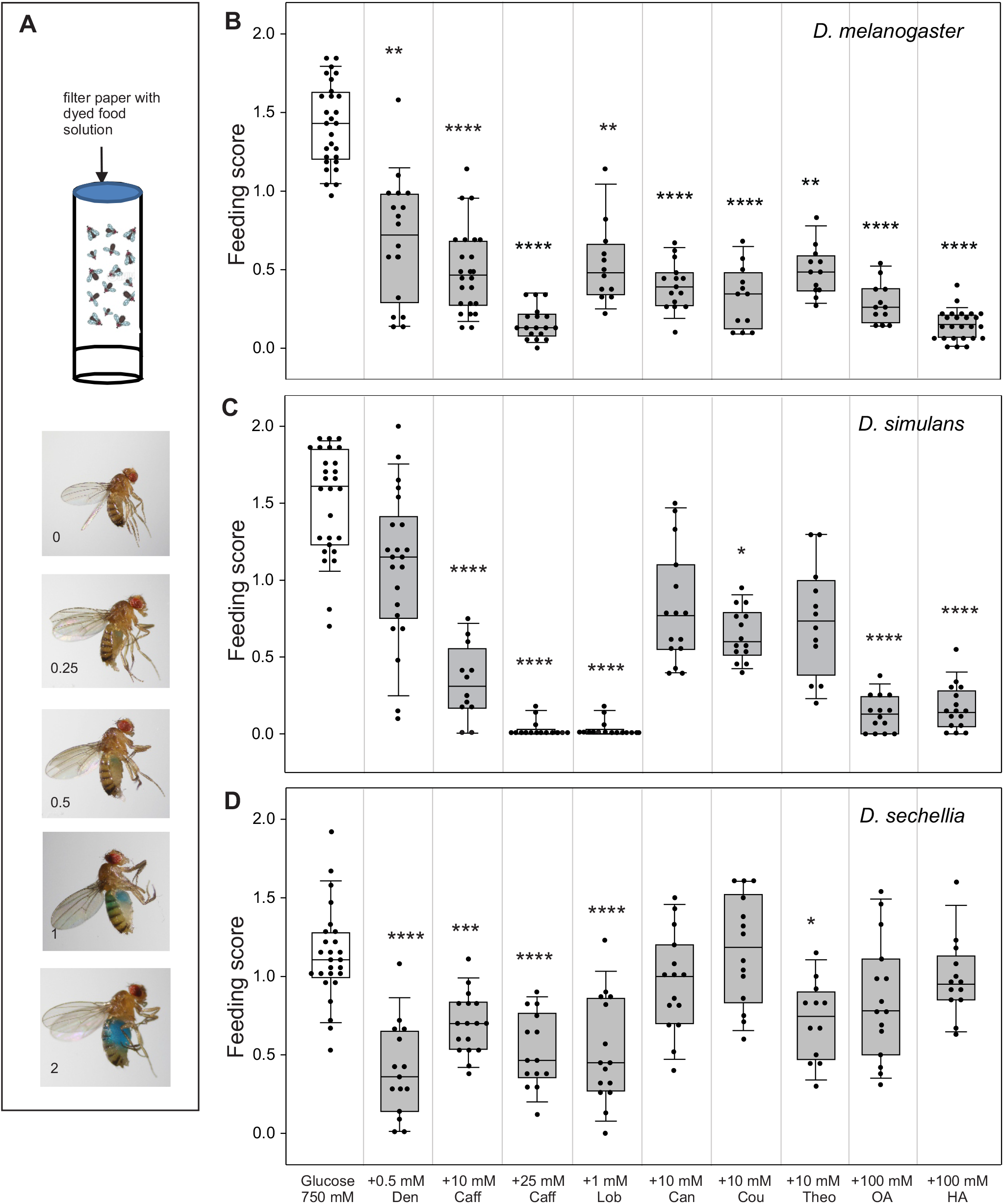
Bitter compounds evoked different levels of feeding aversion across species. **A.** Schematic representation of the group feeding assay. Flies were food-deprived for 24 hs (n=12-15/vial) and then transferred to vials containing a disk of filter paper impregnated with 160 μl of 750 mM glucose (control vials), or 750 mM glucose + a bitter or fatty acid compound (test vials) dyed blue. After 30 minutes vials were frozen and then flies in each vial were scored blind to treatment according to the amount of blue dye in their abdomen using a five-point scale (0-2) (as in Reisenman et al. 2019); a single feeding score was calculated for each vial (=biological replicate). **B-D:** Feeding scores of female *D. melanogaster*, *D. simulans* and *D. sechellia* offered control (white boxes) or test (gray boxes) food solutions. Test vials had 750 mM glucose plus one of the following: 0.5 mM denatonium (den), 10 mM or 25 mM caffeine (caff), 1 mM lobeline (lob), 10 mM L-canavanine (can), 10 mM coumarin (cou), 10 mM theophylline (the), 100 mM octanoic acid (OA, 1.3% vol/vol), or 100 mM hexanoic acid (HA, 1.6% vol/vol). Boxes and horizontal lines within, whiskers, and symbols represent data as explained in Fig. 2E-F. Asterisks indicate differences against the control for each species (Kruskal Wallis ANOVAs and post-hoc Dunn’s tests; **** p<0.001, ***p<0.005, **p<0.01, *p<0.05; n=12-28/species and food solution). In *D. melanogaster* and *D. simulans* **(B-C)**, but not in *D. sechellia* **(D)**, the two fatty acids significantly reduced feeding. In *D. melanogaster* all bitter compounds reduced feeding **(B)**, while caffeine, lobeline and coumarin reduced feeding in *D. simulans* **(C)**. *D. sechellia* consumed similar food amounts in absence or presence of canavanine or coumarin (**D**, p>0.05).

Food solutions consisted of 750 mM glucose alone (control), or 750 mM glucose plus one of the following test compounds (all from Sigma-Aldrich, Saint Louis, Missouri, USA, unless specified): 0.5 mM denatonium benzoate (CAS # 3734-33-6); 10 and 25 mM caffeine (CAS # 58-08-2); 1 mM (L)-lobeline hydrochloride (CAS # 134-63-4); 10 mM L-canavanine (CAS # 543-38-4); 10 mM theophylline (CAS # 58-55-9); 10 mM coumarin (CAS # 91-64-5); 100 mM (1.3% vol/vol) octanoic acid (CAS # 124-07-2); or 100 mM (1.6% vol/vol) hexanoic acid (MP Biomedicals, Solon, OH, USA, CAS # 142-62-1). High concentrations of glucose elicit strong feeding responses, while addition of bitter compounds reduced taste and feeding responses (Dweck and Carlson, 2019; French et al., 2015b; Ling et al., 2014; Weiss et al., 2011). Denatonium is a synthetic compound; lobeline, theophylline and caffeine are plant-produced alkaloids; canavanine is a plant-produced aminoacid; caffeine and canavanine defend plants against herbivory (Fürstenberg-Hägg et al., 2013). Octanoic and hexanoic acid are prominent compounds in *M. citrifolia* ripe fruits (Pino et al., 2010) which are harmless to *D. sechellia*, but toxic (at concentrations >1%) to *D. melanogaster* and *D. simulans* (Hungate et al., 2013; Jones, 1998). We also used pure Noni juice dyed blue. Noni contains anthraquinones and the coumarin-derivative scopoletin (Deng et al., 2010; Ikeda et al., 2009; Satwadhar et al., 2011; Singh, 2012), both of which are insoluble in water, precluding testing. In two control experiments, flies were allowed to feed on 750 mM glucose or water dyed blue in the presence of octanoic acid vapors (10 μl of 100 mM solution or the mineral oil solvent loaded in filter paper), as described in (Reisenman and Scott, 2019) (Figure S3A). As much as possible, food solutions were tested in parallel with overlapping cohorts of flies. At least one control test (750 mM glucose alone) for each species was always conducted along with tests with bitter/fatty acid compounds (i.e. at the same time and with overlapping fly cohorts). Because excess control data-points were therefore obtained over the course of all experiments, control data for each species/genotype was randomly eliminated to achieve sample sizes comparable to those of flies tested with bitter/fatty acid compounds.

Raw and normalized feeding scores were respectively used for comparisons within or between species and sexes. Normalization also served to account for differences in basal (glucose) consumption between strains and genotypes, and for potential differences in ingestion which could result from variations in room temperature, age, and fly cohort. Normalized feeding scores were calculated as: feeding score (glucose + bitter/fatty acid) / feeding score (glucose only). Feeding scores from vials with control data (i.e. glucose only) obtained in the same day and species were averaged and this average was used for normalization. Thus, normalized values circa 1 indicate no difference in consumption between control and solutions containing a test compound, and values <1 and >1 indicate feeding aversion and enhancement, respectively. To established the behavioral valence of each compound, normalized feeding scores were compared against the expected median=1 (no differences in consumption between test and control solutions) using one-sample signed rank tests (Zar, 1999).

Mann-Whitney U tests were used to compare two independent groups. Kruskal-Wallis ANOVAs were used for comparing more than two groups; significant results were followed by Dunnett (for comparisons involving equal sample sizes) or Dunn’s tests (for comparisons involving unequal sample sizes or to compare control vs. all experimental groups) (Zar, 1999).

## RESULTS

### Taste responses are organ-specific

PER to highly appetitive stimuli, 1 M glucose or sucrose, are organ-dependent: tarsi stimulation evoked stronger responses than proboscis stimulation in *D. sechellia* but not in *D. melanogaster* (Fig. 1A-B), and this pattern was consistent across *D. sechellia* strains and sexes (Fig. S1 B-C). At lower sugar concentrations, *D. melanogaster* showed higher PER upon proboscis stimulation (Fig. S1C). In both species, PER to 1 M sucrose was strongly reduced upon addition of the bitter compound denatonium (Fig. 1C). These results demonstrate organ-specific appetitive and aversive taste responses across species.

### *D. sechellia* has increased taste and feeding responses to Noni

Individual flies were first tarsi-stimulated with 1 M sugar, offered water, and then tarsi-stimulated three times with Noni reconstitute. *D. simulans* and *D. melanogaster* had much higher PER rate to stimulation with sugar than with Noni (90-100% and 25-40% respectively, Mc Nemar tests, p<0.0001 for both species), while *D. sechellia* had similar responses to both stimuli (73 and 63% PER, p>0.05; Fig. 2A). To probe whether reduced PER to Noni stimulation in the generalists was due to olfaction, we tested 24 hours food-deprived intact *D. melanogaster* or with their olfactory organs (antennae and maxillary palps) removed two days before experiments. Both groups of flies had indistinguishable PER to Sucrose and Noni, as in Fig. 2A, confirming that the aversion to Noni is solely due to taste (Fig. 2B; Brown et al. 2021).

*D. melanogaster* and *D. sechellia* individual flies were also assayed in a temporal consumption assay. Flies were prepared as before (Fig. 1A), and their tarsi were repeatedly stimulated with 750 mM glucose or Noni juice and allowed to drink. A similar proportion of flies from each species fed on glucose (81% *D. melanogaster* and 91% *D. sechellia;* Fisher Exact test, p>0.05; n=21, 22), but a larger proportion of *D. sechellia* flies fed on Noni (31.6% and 78.3%, respectively; p<0.0074, n=19, 23; Fig. 2F). *D. melanogaster* consumed for longer and fed more times upon stimulation with glucose than with Noni, but these behavioral metrics were not different in *D. sechellia* (Fig. 2D-E). Thus, Noni is comparatively more appetitive in *D. sechellia* than in *D. melanogaster*.

### *D. sechellia* has reduced feeding aversion to bitter compounds

We then switched to a group feeding assay where flies were unrestrained (Fig. 3A) to investigate responses to known bitter stimuli or Noni fatty acids across species. Flies were offered 750 mM glucose (control), or 750 mM glucose + a test stimulus (bitter or fatty acid). Addition of any of the test stimuli significantly reduced feeding in *D. melanogaster* (Fig. 3B, Kruskal-Wallis ANOVA followed by Dunn’s tests, p<0.005); *D. simulans* had reduced feeding on solutions containing caffeine, lobeline, coumarin, or the two fatty acids (post-hoc Dunn’s tests, p<0.05; Fig. 3C). In *D. sechellia* addition of canavanine, coumarin, or either of the fatty acids no impact (post-hoc tests, p>0.05; Fig. 3C). To evaluate potential lethal effects of octanoic acid in the generalist species, individual flies were placed in vials offering 750 mM glucose alone or with octanoic acid, allowed to feed during 30 minutes, and we recorded viability. None of the flies offered only glucose died, but 4/16 and 1/19 *D. melanogaster* and *D. simulans* respectively were dead or stuck to the food, but otherwise flies seemed normally active. The remaining 12 and 18 flies of each species fed significantly less on solutions with octanoic acid (Fig. S2). We also tested whether feeding aversion to octanoic acid could be due to olfaction. *D. melanogaster* flies were offered 750 mM glucose in presence or absence of octanoic acid odors (no contact, as illustrated in Fig. S3A), and found that flies from both groups fed indistinguishable large amounts of glucose (Fig. S3B). Overall, these results confirmed that the aversion to octanoic acid is mediated solely by taste and it is not due to lethality.

Feeding on 750 mM glucose (control) varied across species, with *D. sechellia* feeding less than its generalist relatives (Kruskal Wallis ANOVA followed by post-hoc Dunn tests, p<0.01; n=26-29). Thus, for comparing responses across species, we normalized the scores of flies fed test solutions (i.e. 750 mM glucose + bitter/fatty acid) to that of flies fed 750 mM glucose only. This normalization, here and thereafter, additionally accounts for possible cohort, temperature, and day-to-day variability. The normalized feeding scores of the generalist species were significantly <1 for most test compounds (Fig. 4; one sample signed rank tests, p<0.05 in all cases except *D. simulans* tested with denatonium and lobeline). In contrast, the normalized feeding scores of *D. sechellia* fed solutions containing canavanine, coumarin, octanoic acid, and hexanoic acid were not statistically different from 1, indicating no feeding aversion neither appetite (p>0.05 in all cases). *D. sechellia* showed significant feeding aversion to caffeine, lobeline, theophylline, and denatonium (medians<1, p<0.05, Fig. 4). However, for the most part, their normalized scores were divergent from those of *D. melanogaster* and *D. simulans*: responses to canavanine, caffeine, theophylline, and the two fatty acids were similar in the two generalists (Kruskal Wallis ANOVAs followed by Dunn tests, p>0.05) but different from those of *D. sechellia* (post-hoc Dunn tests, p<0.05). In all cases (except denatonium and lobeline) the responses of *D. sechellia* were different from those of *D. melanogaster*. These results were consistent between sexes and for another strain of *D. sechellia* (Fig. S4A-B), showing that the reduced feeding aversion is species- and not strain-specific. Notably, males had feeding aversion to hexanoic acid (Fig. S4A), suggesting that this Noni compound has sex-specific functions (Álvarez-Ocaña et al., 2022). We compared responses to caffeine, an alkaloid in a plant family (Rubiaceae) that includes coffee and Noni (Singh, 2012), and to denatonium (an aversive synthetic compound) across two strains for each species. Responses within a species were consistent, and caffeine evoked similarly higher levels of feeding aversion in the generalists (Fig. S4C). Overall, these results indicate that *D. sechellia* has a lineage-specific reduced sensitivity to canavanine, caffeine, theophylline, coumarin, and to octanoic and hexanoic acids. These are likely a subset of the compounds to which *D. sechellia* has reduced aversion (Dweck and Carlson, 2020).

**Figure 4:**
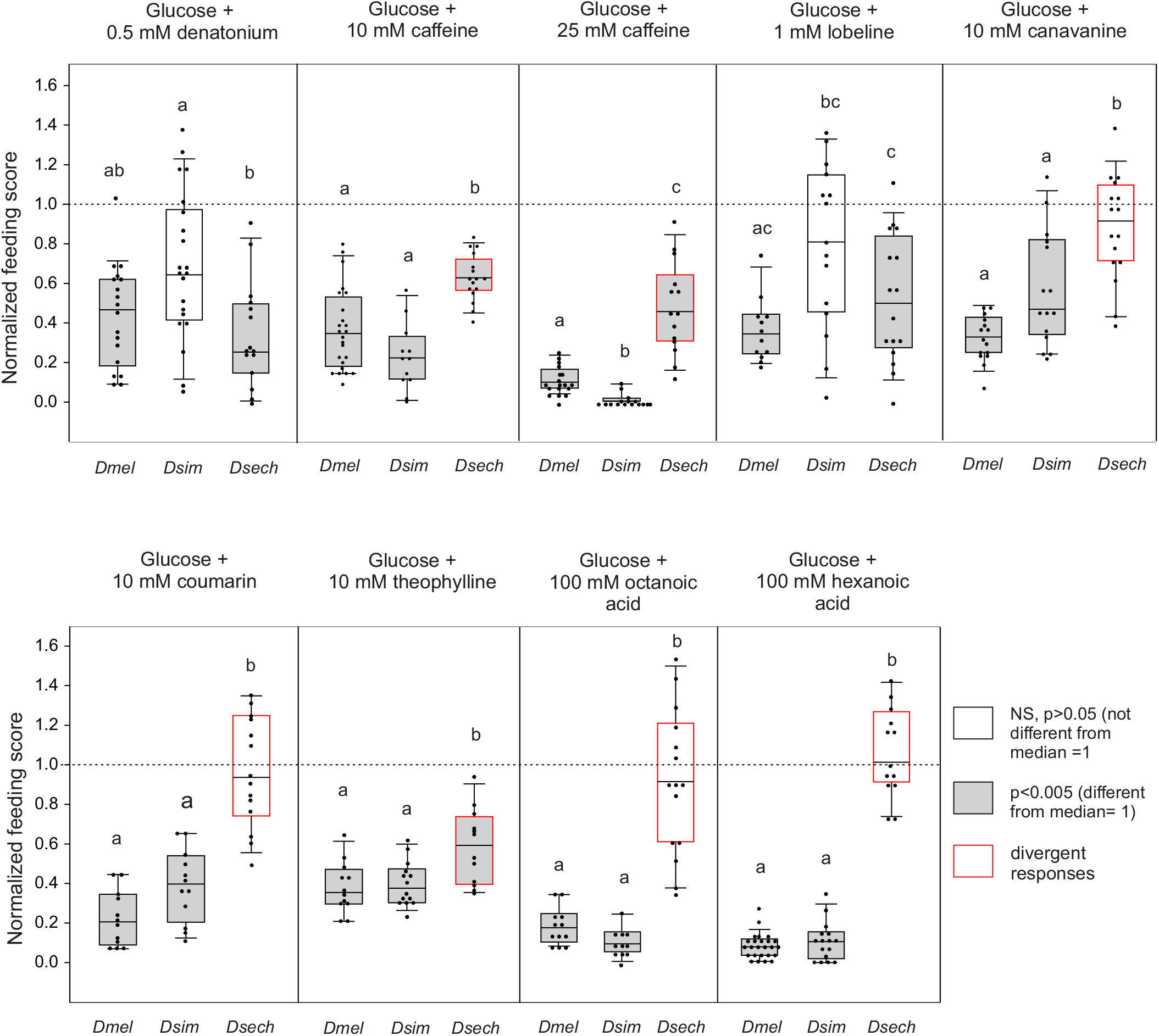
*D. sechellia* has a comparatively reduced feeding aversion to most bitter compounds and Noni fatty acids. Data represent the glucose-normalized (on a day-to-day basis) feeding scores of female *D. melanogaster*, *D. simulans* and *D. sechellia* flies (calculated from data in Figure 3), to allow interspecific comparisons. Data is represented as in Figure 2E-F; n=12-25 species/test compound. The horizontal dotted lines at 1 indicate similar consumption on control (glucose only) and test solutions (glucose + bitter/fatty acid compound), i.e no feeding aversion neither enhancement to the test solutions. *D. melanogaster* and *D. simulans* showed feeding aversion to solutions containing either canavanine, coumarin, octanoic acid, or hexanoic acid (gray shaded bars indicate p-values, one-sample signed rank tests against median=1), while *D. sechellia* lost the aversion to these compounds (white bars, p>0.05 in all cases). *D. sechellia* flies retained aversion (feeding scores significantly<1) to caffeine, lobeline, theophylline and denatonium (gray shaded bars). In all cases except for lobeline and denatonium, the responses of *D. sechellia* flies were divergent from those of *D. simulans* and *D. melanogaster* (boxes outlined red, p<0.05 in all cases; different letters indicate inter-specific differences, Kruskal Wallis ANOVAs and post-hoc Dunn’s tests).

### Two GRs losses correlate with the reduction of aversive responses to bitter compounds

We next investigated whether the reduced aversion to bitter compounds in *D. sechellia* correlates with the losses of GRs in this species. McBride (2007) showed that at least 14 *GRs*, several of which mediate responses to bitter compounds in *D. melanogaster*, are lost or pseudogenized (Table 1). We focused on *GR28b.a* and *GR39a.a*, as these are widely expressed (75-100%) in *D. melanogaster* bitter sensilla (Ling et al., 2014; Weiss et al., 2011; Table 1). Albeit *GR39a* and *GR28b* undergo alternative splicing in drosophilid flies (Gardiner et al., 2008; Sang et al., 2019), we used null *D. melanogaster* mutants for each of these *GR* genes. We hypothesized that one or both mutants would recapitulate the *D. sechellia* lineage-specific reduced aversion to bitter compounds specifically but not to fatty acids, as these are mediated by IRs (Ahn et al., 2017; Masek and Keene, 2013). Feeding responses were normalized as before, and the responses of mutants were compared to those of their respective genetic controls (rather than to those of wildtype flies), as it is standard in the field. The normalized feeding scores of *D. melanogaster* flies from all genotypes (mutants and genetic controls) offered any of the test compounds (except GR39a mutants offered canavanine) were significantly <1 (Fig. 5, one-sample signed rank tests, p<0.05), indicating reduced feeding for test solutions containing any of the test chemicals, but not loss of significant aversion as observed for some compounds in *D. sechellia*. *GR28b* and *GR39a* mutants, however, fed significantly more than their respective genetic controls on solutions containing canavanine or coumarin, and *GR28b* mutants fed more than its control on solutions containing theophylline (Mann-Whitney U tests, p<0.05 in all cases). In addition, mutants showed a generalized reduction of aversion to bitter compounds (e.g. responses to caffeine and theophylline in *GR39a* mutants were near-significance, p=0.066 and p=0.051 respectively, consistent with Dweck and Carlson, 2020). As expected, aversion to fatty acids remained unaltered in mutants (p>0.05). Overall, these results demonstrate that losses of single bitter *GRs* are sufficient to reduce feeding aversion. In particular, the losses of two *GRs* which are widely expressed in bitter GRNs in *D. melanogaster* collectively correlate with the *D. sechellia’s* loss of aversion to canavanine, theophylline, and coumarin.

**Table 1:**
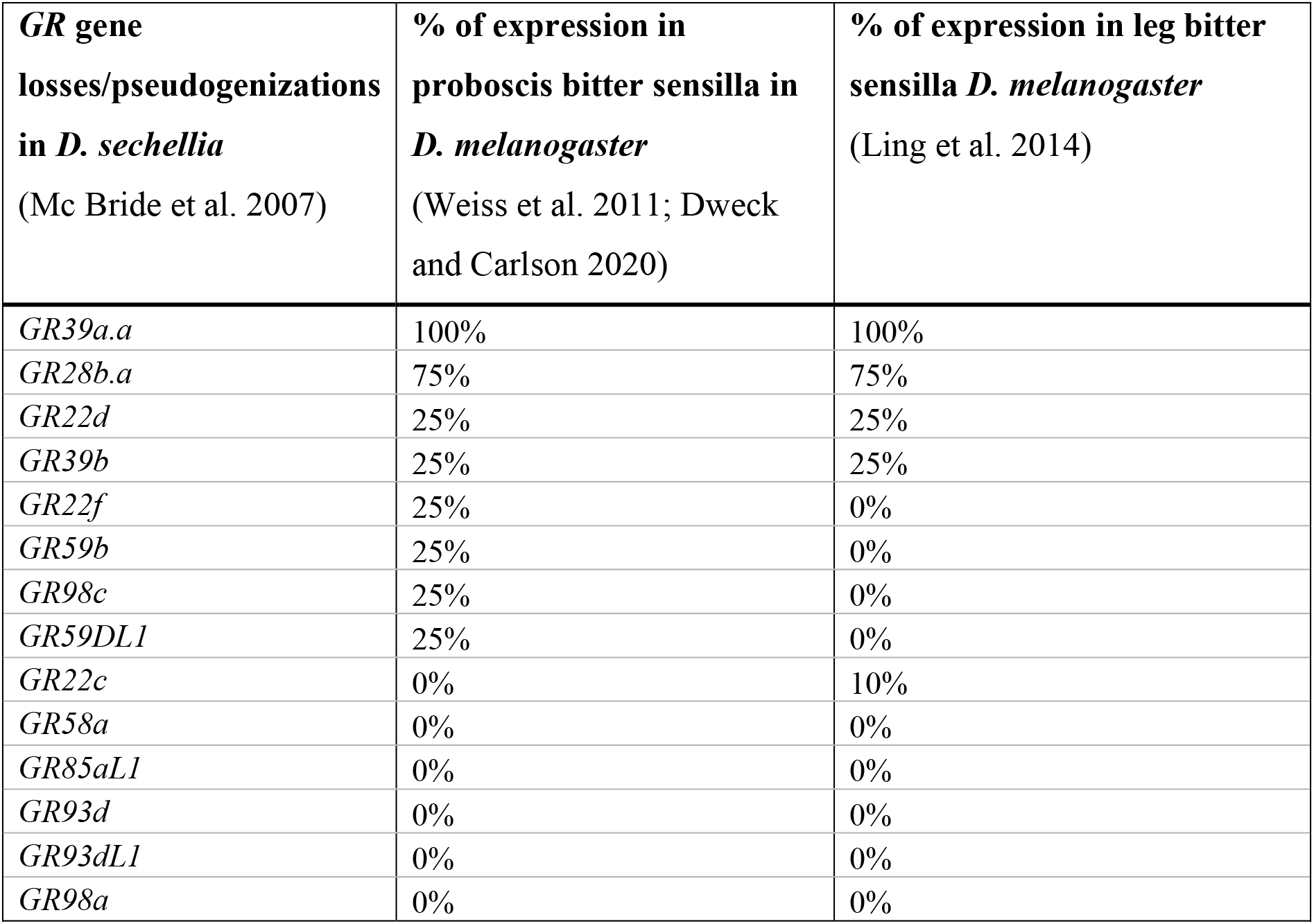
Gustatory receptor losses in *D. sechellia* (McBride, 2007) and percentage of expression in bitter GRNs of the proboscis and legs of *D. melanogaster* (Ling et al., 2014; Weiss et al., 2011). *GRs* are in descendent order of expression in *D. melanogaster*. GR39a.a and GR28b.a are widely expressed in *D. melanogaster* and lost in *D. sechellia* (and also in the specialist *D. erecta);* GR39a.a is important for detection of many bitter substances in *D. melanogaster* (Dweck and Carlson, 2020).

**Figure 5:**
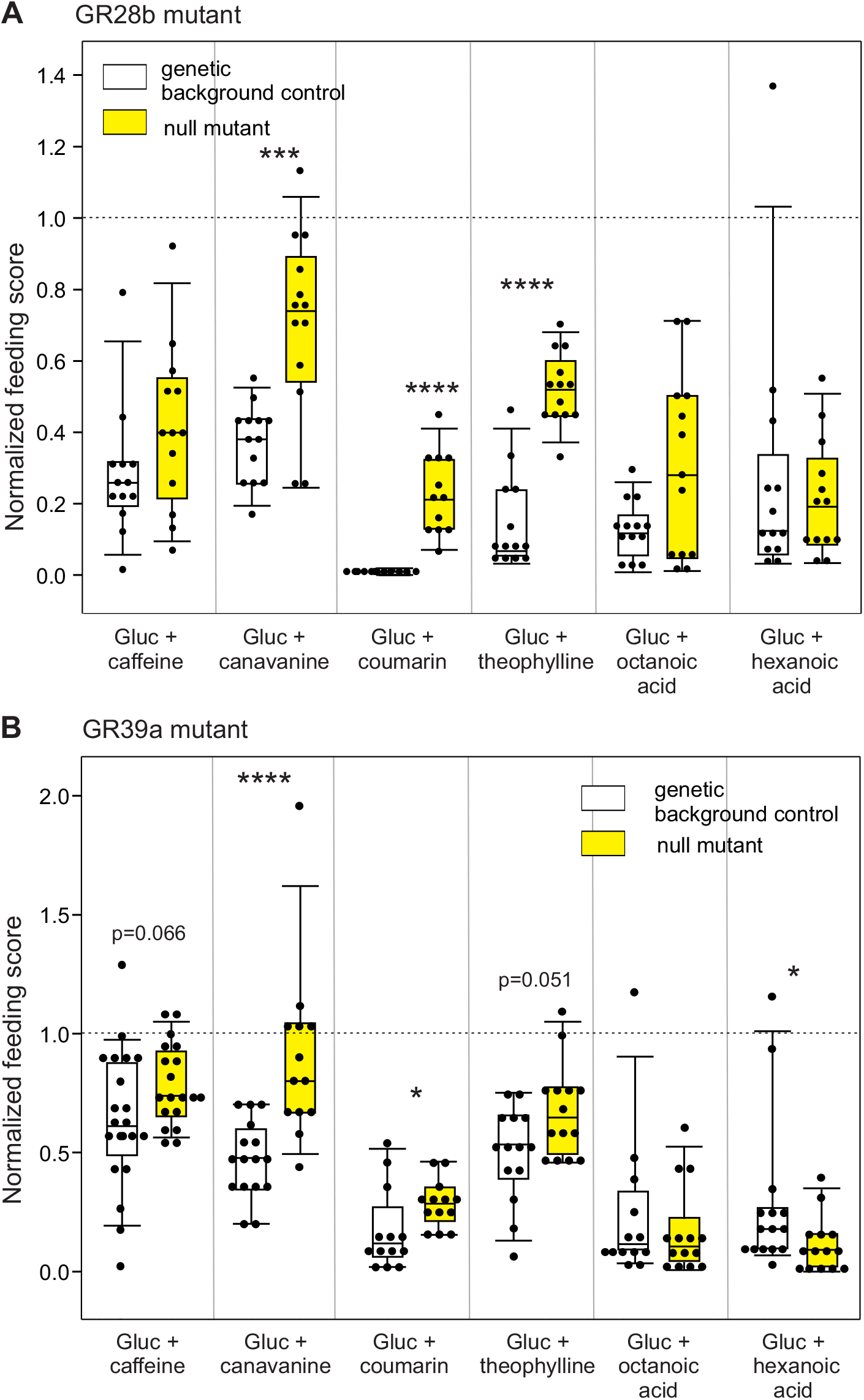
GR28b and GR39a *D. melanogaster* null mutants have reduced aversion to bitter compounds that correlates with the *D. sechellia’s* behavioral phenotype. Glucose-normalized feeding scores (calculated as in Figure 4) of GR28b (**A**) and GR39a (**B**) null mutants (yellow boxes) and their respective genetic background controls (white boxes) to solutions containing 750 mM glucose plus a bitter or fatty acid compound (concentrations as in Figs. 3–4). Data representation as in Fig. 2E-F; the horizontal dotted lines at 1 indicate no feeding aversion neither enhancement. Addition of any bitter/fatty acid compound reduced feeding in all (one-sample signed rank tests, p<0.005) but one case (GR39a mutants offered canavanine; p>0.05); n=12-21 for each genotype/food solution. Both mutants consumed larger amounts of solutions containing canavanine or coumarin (and theophylline in GR28b mutants) than their respective genetic controls (**A**, Mann-Whitney U tests, asterisks, ***p<0.005, ****p<0.001); aversive responses to caffeine and theophylline were slightly reduced in GR39a mutants (**B**, p=0.066 and 0.051, respectively). Responses to fatty acids were not different between mutants and their respective controls for the most part (Mann-Whitney U test, p>0.05; GR39a mutants consumed less than its control, *p<0.05).

### Octanoic acid enhances feeding in *D. sechellia*

Addition of fatty acids to appetitive solutions significantly reduced feeding in the generalist species but not in *D. sechellia* (Figs. 3–4), suggesting that proteins that detect them may be lost (or have lower expression) in bitter GRNs (Ahn et al., 2017). In *D. melanogaster*, concentrations <1% vol/vol are non-toxic and stimulate sugar GRNs (GR64e-positive, Kim et al., 2018), evoking appetitive responses. We thus investigated if octanoic acid at >1% (octanoic acid is highly concentrated in Noni) can increase feeding in *D. sechellia*, rather than aversion. We tested parallel cohorts of flies with 750 mM glucose, water only, or water + 100 mM (1.3% vol/vol) octanoic acid. As expected, in both species, the feeding scores of flies fed glucose were much higher than those of flies offered water or water + octanoic acid (Fig. 6A, Mann-Whitey U tests followed by Dunn’s tests, p<0.05). Glucose responses normalized to water were >1 in both species (one-sample Signed rank tests, p<0.01 in both cases; Fig. 6B, left), indicating feeding enhancement. The water-normalized scores of *D. melanogaster* to octanoic acid were aversive (<1; p<0.001), while those of *D. sechellia* were appetitive (>1, p<0.01; Fig. 6B, right). The water-normalized responses of *D. sechellia* to glucose or octanoic acid were higher than those of *D. melanogaster* (Mann-Whitney U tests, p<0.05; Fig. 6B). In addition, *D. sechellia* consumed similar amounts of water in presence or absence of octanoic acid odors (no contact allowed as illustrated in Fig. S3A; Mann-Whitney U test, p>0.05; Fig. S6), which indicates that olfaction alone is not sufficient to increase consumption (Masek and Keene, 2013). This shows that octanoic acid, a prominent toxic host-compound, increases feeding in *D. sechellia* at concentrations that are aversive to its generalist relatives.

**Figure 6:**
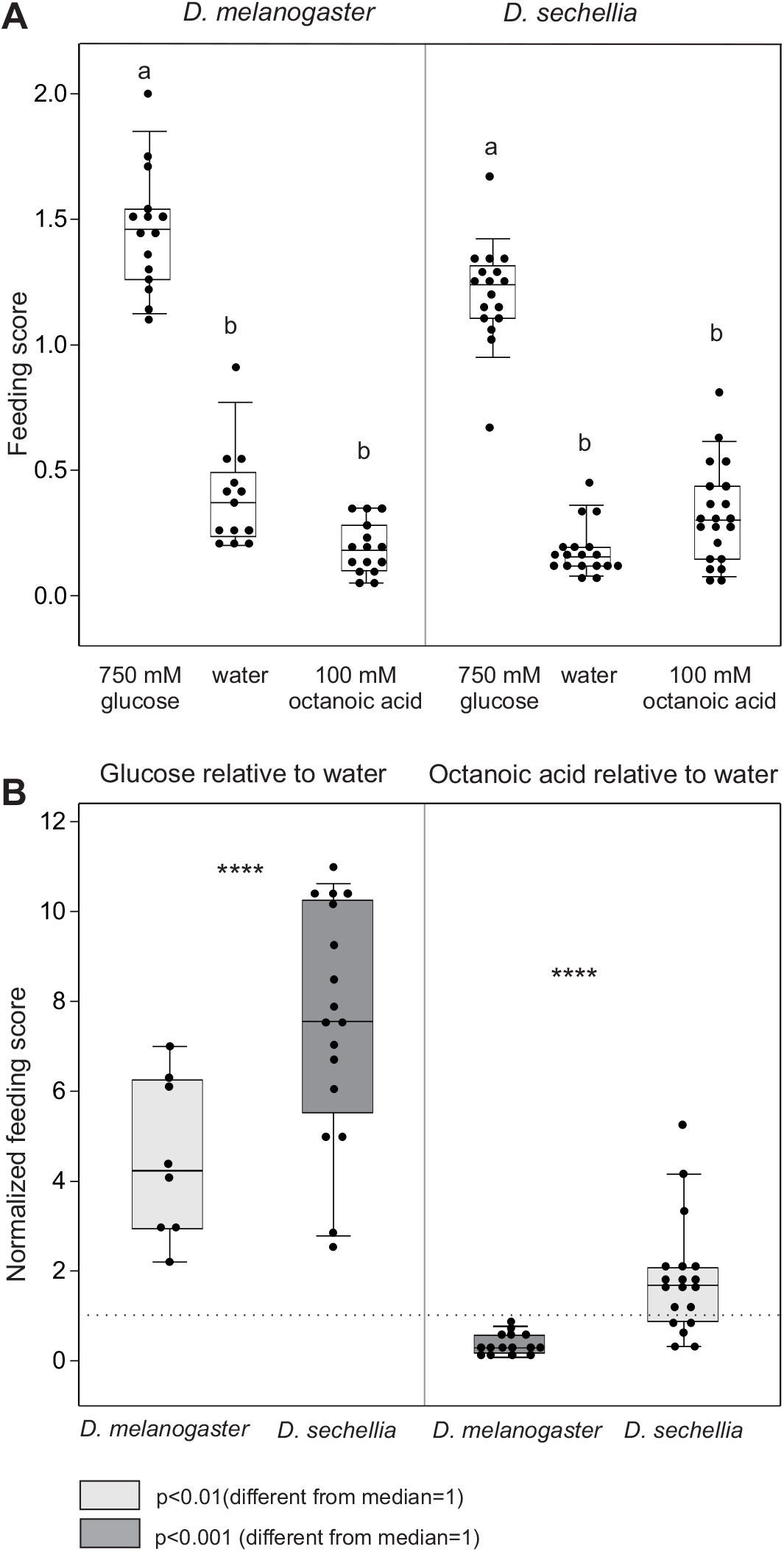
*D. sechellia* has increased appetite for Noni fatty acids in comparison with *D. melanogaster*. **A:** Feeding scores of *D. melanogaster* (left) and *D. sechellia* (right) offered 750 mM glucose, water, or water + 100 mM (1.3% vol/vol) octanoic acid. Flies were prepared as before, and all groups were tested with parallel cohorts. Both species fed the most on glucose (Kruskal-Wallis ANOVAs followed by Dunn’s tests, n=13-21/species and food solution; different letters indicate significant differences, p<0.05). **B:** Water-normalized feeding scores of flies offered glucose (left) or octanoic acid (right). The horizontal dotted line at 1 indicates no aversion neither preference in comparison to water; gray shades indicate p-values (one-sample signed rank tests against median=1). Relative to water, both species have strong appetite for glucose (i.e. median>1; left), but *D. melanogaster* fed less on octanoic acid (i.e. median<1), while *D. sechellia* fed more (i.e. median >1, right). Normalized responses to glucose and octanoic acid differed between species (Mann-Whitney U tests, asterisks, ***p<0.005, ****p<0.001).

## DISCUSSION

Chemosensory specializations allow specialist insects to find their hosts. Host-olfactory adaptations are more understood than taste adaptations, and include a gain and/or increase in peripheral sensitivity to host odors. To investigate taste adaptations, we took advantage of the specialist *D. sechellia*, which is closely related to the generalists *D. simulans* and *D. melanogaster*. *D. sechellia* feeds and oviposits almost exclusively on ripe Noni fruit, which is toxic (due to its high content of fatty acids) to many drosophilid species. Using taste and feeding assays, we found that in comparison with its close relatives, adult *D. sechellia* has reduced feeding aversion to various bitter substances that correlate with the lineage-specific losses of various *GR* genes, including the broadly expressed *GR39a.a* and *GR28b.a*. Furthermore, *D. sechellia* has increased appetite for octanoic acid, a signature Noni fatty acid compound. Our studies powerfully demonstrate the role of GRs in mediating taste specialization in *D. sechellia*.

### Appetitive responses are organ-specific in *Drosophila sechellia*

Using PER, which measures taste-detection in the absence of feeding, we found species- and organ-specific differences: *D. sechellia* had stronger PER to sugar stimulation of the tarsi than to proboscis stimulation, while *D. melanogaster* showed similar responses (Figs. 1). Moreover, at low concentrations, proboscis stimulation evoked higher PER in *D. melanogaster* (Fig. S1), consistent with the highest responsiveness of proboscis taste sensilla in this species (Dahanukar et al., 2007; Ling et al., 2014). Because the numbers and positions taste sensilla in the proboscis are similar between the two species (Dweck and Carlson, 2020), species might differ in the expression of conserved sweet *GRs* (e.g. *GR5a*, *GR61a* and *GR64a-f*) in this organ (Dahanukar et al., 2007; Freeman et al., 2014). Differences in GRN projections can also contribute to these differences: most tarsi GRNs terminate in the ventral nerve cord, while all proboscis GRNs terminate in the brain primary taste center. Furthermore, different circuits control distinct phases of the feeding behavioral program, such as PER versus ingestion (Chen and Dahanukar, 2020; Scott, 2018). For example, a subset of sweet tarsal GRNs are necessary for appetitive responses to sugar and another subset for stopping locomotion upon food encounter (Thoma et al., 2016). This functional subdivision may be relevant for specialists, as the legs are the first appendages that contact potential food sources. PER was inhibited by the synthetic bitter compound denatonium in both *D. melanogaster* and *D. sechellia* (Fig. 1C), revealing a fundamental principle for avoiding ingestion of potentially toxic substances.

### *D. sechellia* has increased taste and feeding responses to Noni

Although *D. sechellia* is found preferentially on Noni, flies have been reported in the Seychelles from mangoes and figs (Matute and Ayroles, 2014). *D. sechellia* uses olfactory cues for host orientation and finding (Auer et al., 2020, 2021; Zhao and McBride, 2020), but it is not clear whether taste cues play a distinct role for food acceptance, although females require both taste and olfactory input for oviposition on Noni (Álvarez-Ocaña et al., 2022). Noni fruit is difficult to source, and therefore we used dehydrated fruit leather strips and juice. The juice has a consistent composition and has been used in chemical ecology studies (Auer et al., 2020; Álvarez-Ocaña et al., 2022), although it has altered amounts of some compounds (Abou Assi et al., 2017; Almeida et al., 2019; Auer et al., 2020; Motshakeri and Ghazali, 2015). Nevertheless, we found that *D. sechellia* has similar taste and feeding responses to high sugar and Noni products, while the responses of the two generalists to Noni were much reduced (Fig. 2). Furthermore, the taste and feeding aversion to Noni and octanoic acid in the generalists is not due olfaction (Fig. 2B and S3).

### *D. sechellia* has reduced feeding aversion to various bitter compounds

A common theme among drosophilid and other insects with specialized diets is the loss of taste chemoreceptors for detection of bitter (likely noxious) compounds (Crava et al., 2016; Dweck and Carlson, 2020; Dweck et al., 2021; McBride, 2007; Rytz et al., 2013; see Briscoe et al., 2013; Wanner and Robertson, 2008 for exceptions). This is because specialists developed resistance to host toxins, providing a private ecological niche, but also because they do not longer encounter them due to their specialization in one or a few related hosts. Generalist insects, in contrast, use many food sources that might contain diverse potentially noxious (bitter) substances, which need to be detected and their ingestion prevented. In particular, *D. sechellia* lost various *GRs* and divergent *IRs* (Crava et al., 2016; McBride, 2007; McBride and Arguello, 2007; Rytz et al., 2013), including *GR39.a.A*, which is expressed in all bitter GRNs in *D. melanogaster* and is important for the detection of various bitter compounds in adults (Table 1, Dweck and Carlson, 2020; Ling et al., 2014; Weiss et al., 2011) and larvae (Choi et al., 2016; Choi et al., 2020; Kwon et al., 2011). Our behavioral results are in line with these predictions: *D. sechellia* have reduced, and even abolished, aversive feeding responses to compounds that suppress consumption in the generalist ancestors, most notably *D. melanogaster* (Figs. 3–4; Fig. S4). Moreover, the responses of *D. sechellia* to caffeine, canavanine, coumarin and theophylline were different from those of *D. simulans* and *D. melanogaster* (Fig. 4). *D. sechellia* retained aversive responses to lobeline, a toxic plant alkaloid which decreases feeding (Alsop, 2012; Wink, 1998) and to the synthetic compound denatonium, which is unpalatable but may not be toxic. Studies from labellar bitter-sensitive sensilla in *D. sechellia* also found no responses to theophylline and caffeine but strong responses to lobeline, denatonium, and surprisingly, also to coumarin (Dweck and Carlson, 2020). Coumarin is a precursor of the signature Noni compound scopoletin (Ikeda et al., 2009), and thus it is possible that this compound has different effects depending on the behavioral context. For instance, GR66a-positive GRNs mediate behaviors of opposite valence such as positional aversion and oviposition attraction, dependent on the taste organ (Joseph and Heberlein, 2012). Overall, our results are in line with the prediction that specialist insects lost aversion to bitter compounds in general because these are no longer encountered and/or are important for host acceptance. Reduction of bitter sensitivity was reported in non-specialists: the generalist *D. suzukii* is a pest of thin-skin fruits (e.g. berries, cherries) and has reduced sensitivity to e.g. caffeine and theophylline, allowing flies using the un-ripen fruit stages which contain large amounts of bitter substances (Durkin et al., 2021; Dweck et al., 2021; Karageorgi et al., 2017).

### *D. sechellia* lost the feeding aversion to Noni fatty acid compounds

The fatty acid compounds hexanoic and octanoic acid are main Noni compounds (19% and 58% respectively; Farine et al., 1996), and are toxic to many drosophilids including *D. melanogaster* and *D. simulans*, but not to *D. sechellia* (Legal et al., 1994). We found that *D. sechellia* lost the ancestral taste aversion to hexanoic and octanoic acid (Figs. 3–4; Fig. S4). In *D. melanogaster*, aversion to >1% vol/vol of fatty acids involves activation of tarsal bitter GRNs labeled by GR33a (Ahn et al., 2017; Chen and Dahanukar, 2020; Prieto-Godino et al., 2017), but the specific GRs necessary for this aversion remain uncharacterized. A study in a population of *D. yacuba* which uses Noni similarly to *D. sechellia* identified selection in four bitter *GR* genes (*GR22b*, *GR22d*, *GR59a* and *GR93c*), but neutral evolution in *IR* genes mediating appetite to low fatty concentrations via sweet GRNs (Ahn et al., 2017; Ferreira et al., 2020; Yassin et al., 2016). In *D. sechellia* various GRs of the GR22a clade have been pseudogenized (McBride, 2007; Table 1). Overall, these results suggest that the *D. sechellia’s* lack of aversion to fatty acids at >1% vol/vol (Figs. 3,4) maybe due to evolutionary changes in certain chemosensory proteins others than GRs expressed in bitter GRNs, such as losses/pseudogenizations and/or qualitative and quantitative changes in the combinations of proteins expressed therein.

Another noteworthy finding is that *D. sechellia* males have aversion to hexanoic (but not to octanoic) acid, and consumed much less on these solutions than females (Fig. S4A). This suggests a female-specific role for this compound, in line with the findings that hexanoic acid is a more efficient oviposition attractant than octanoic acid (Amlou et al., 1998; Álvarez-Ocaña et al., 2022). Furthermore, oviposition preference for hexanoic acid requires both olfaction (via IR75a) and taste (Álvarez-Ocaña et al., 2022). Female *D. sechellia* intently probes the substrate with its ovipositor (Álvarez-Ocaña et al., 2022), but it is unclear whether this serves to evaluate substrate chemical composition. The female terminalia of *D. melanogaster* has trichoid sensilla (Stocker 1994, Taylor 1989) and expresses various GRs, IRs and OBPs (Crava et al. 2020), but its chemosensory function has not been proven. Alternatively, the loss of feeding aversion to hexanoic acid and the oviposition preference for this compound might result from differential expression of IRs in tarsi, similar to the IR76b/IR25a-mediated *D. melanogaster’s* female preference for acidic substrates (Chen and Amrein, 2017). Sexual dimorphism in taste chemosensory proteins mediating oviposition in host plants occurs in the specialist butterfly *Heliconius melpomene* (Briscoe et al., 2013), suggesting commonalities in specialists from diverse insect orders.

Albeit toxic at high concentrations, fatty acids are caloric and in *D. melanogaster* evoke appetitive responses (at <1% vol/vol) that require IR56d in sweet (GR64f-positive) GRNs (Masek and Keene, 2013; Tauber et al., 2017). *D. sechellia* not only lost the taste-mediated aversion to solutions containing 1.3% vol/vol (100 mM) octanoic acid (Figs. 3–4; Fig. S4A-B), but had increased appetite for this compound (Fig. 6). *D. melanogaster* had significant aversion to 1.3% vol/vol octanoic acid but *D. sechellia* had increased feeding (Fig. 6B). The enhanced feeding of *D. sechellia* was not due to smell (Fig. S6), similarly to the *D. melanogaster* persistent appetite for low concentrations of fatty acids in absence of olfactory input (Masek and Keene, 2013). It is possible that the *D. sechellia’s* observed appetitive for higher concentrations of fatty acids also involves IR56a/d and IR76b in sweet GRNs (Ahn et al., 2017). Future studies addressing the expression of the chemosensory proteins, in particular IRs, will illuminate the cellular mechanisms mediating taste responses to these important hostplant compounds.

### *D. sechellia’s* reduced food aversion to bitter compounds correlates with losses of single GRs

In the last decade, genomes for many insect species have allowed making predictions about the genomic bases of chemosensory adaptation (see e.g. Vertacnik and Linnen, 2017 and Robertson, 2019, for reviews on this topic), but functional studies to test such predictions, particularly in the case of taste, are more scarce. In *D. sechellia* various hypotheses derived from genomic studies have been tested: changes in specific ORs and IRs mediate olfactory preference for Noni (Álvarez-Ocaña et al., 2022; Auer et al., 2020, 2021), and the losses of two *OBPs* enabled contact-mediated acceptance of fatty-acid oviposition substrates (Matsuo et al., 2007). *D. sechellia* also lost many GRs and various divergent IRs (McBride, 2007), but the functional consequences of these losses remain mostly unexplored. We found that the reported losses of *GR39a.a* and *GR28b.a* correlate with the *D. sechellia’s* behavioral phenotype. *D. melanogaster* null mutants for each of those genes lost the feeding aversion for canavanine and coumarin (and have reduced caffeine aversion), similarly to *D. sechellia’s* (Fig. 5). Although we used null mutants in these experiments, GR39a.a is the splice form of GR39a that mediates responses to various bitter compounds (Dweck and Carlson, 2020), and GR28b, GR28b.c and GR28b.d are respectively involved in saponin detection and temperature sensing (Ni et al., 2013; Sang et al., 2019). As expected, both null mutants retained the aversion to fatty acids, as their detection may involve divergent IRs (Ahn et al., 2017; Brown et al., 2021; Masek and Keene, 2013; Sánchez-Alcañiz et al., 2018). Interestingly, the *GR39a.a* isoform has a large variation in copy number across various drosophilid species examined (Gardiner et al., 2008), and *GR39a.a* and *GR28b.a* are also lost in the specialist *D. erecta* (McBride, 2007). Altogether, these findings suggest that evolution of these *GRs*, whether losses, duplications, and/or changes in expression, are particularly important for the specialization in drosophilid flies.

In *D. melanogaster*, various studies used genetic tools to uncover the GRs that confer responses to bitter compounds. Early investigations showed that one or two GRs were important for responses to bitter substances (e.g. caffeine, Lee et al., 2009; Moon et al., 2006), but then a picture emerged consistent with a model in which multiple GRs act as heteromeric complexes. For instance, GR8a, GR66a and GR98b are important for detection of canavanine (Shim et al., 2015), and co-expression of GR32a, GR59c and GR66a confers sensitivity to lobeline, berberine and denatonium (Sung et al., 2017; reviewed in Chen and Dahanukar, 2020 and Delventhal and Carlson, 2016). Dweck and Carlson (2020) showed that *GR39a.a*, which is lost in *D. sechellia*, is necessary for responses to coumarin and caffeine in *D. melanogaster’s* labellar bitter GRNs, in line with our results.

Studies aimed at discovering the functional consequences of evolution of chemosensory proteins mediating feeding are hindered by the fact that each GRN expresses multiple GRs, and that expression of the same GR in different GRNs produces different responses (Dweck and Carlson, 2020). In addition, individual bitter GRs interact in different ways, providing another strategy for evolution of novel responses (Delventhal and Carlson, 2016). These principles of organization differ greatly from those of the olfactory system, i.e. for the most part a one-to-one cell-olfactory protein, plus one or more co-receptors (Vosshall and Stocker, 2007; Task et al., 2022). Consequently, genomes studies can better guide functional testing of chemosensory proteins mediating olfaction. For instance, changes in one or few ORs and/or IRs can confer new, dramatic adaptive responses in specialist insects (e.g. Álvarez-Ocaña et al., 2022; Auer et al., 2020; Liu et al., 2020; Matsunaga et al., 2022). However, we found that the reported losses of single *GRs* can explain the *D. sechellia’s* phenotype to bitter compounds such as canavanine, theophylline and coumarin (Figs. 3–4; Fig. S4), but it is likely that the many *GR* (and divergent *IR*) gene losses in this species confer, most likely in combination, the complete behavioral phenotype. Although this remains to be investigated, it is expected that larvae, which are immersed in Noni, not only have reduced aversion to host and bitter compounds but have much increased appetite for host fatty acids. The expression of GRs and IRs in *D. sechellia* larvae are not yet characterized, but *D. melanogaster* larvae expresses 39/68 GRs (including GR39a.a and GR28b.a; Kwon et al. 2011, Choi et al. 2016, 2020) and various IRs (Sanchez-Alcaniz 2018, Ni 2020). As observed for adult *D. melanogaster*, their larvae have feeding aversion to e.g. caffeine, canavanine and coumarin, some of which require GR39a (Choi et al. 2016, 2020), highlighting the permissive role of this GR loss in *D. sechellia*. Fatty acids are likely detected by IRs in *D. sechellia* larvae, possibly inducing ingestion depending on their cellular compartmentalization, in line with the finding that *D. melanogaster* larvae has broad expression of IRs (IR25a and IR76b) required for adult fatty acid taste (Chen and Amrein 2017, Ahn et al. 2017; Ni, 2020; Sánchez-Alcañiz, 2018). Future investigations in *D. sechellia* larvae will shed light into the chemosensory adaptations underlying host specialization across life stages.

Finally, in addition to changes in chemosensory proteins and their tissue-specific cellular expression, changes in circuitry, as elegantly reported in the olfactory system of *D. sechellia*, could also play a role in host specialization (Auer et al., 2020). Although such changes tend to be more constrained by pleiotropy (Zhao and McBride, 2020), a recent comparative study proposed that olfactory pathways are conserved, with selection acting in co-expressed copies of cognate ORs (Auer et al., 2021). Future studies, using tools such as CRISPR-Cas9, transcriptomes from various taste tissues/sensilla (e.g. Auer et al., 2020; Dweck et al., 2021), and development of genetically encoded indicators of neural activity in non-melanogaster species, may help addressing these relevant questions.

## Acknowledgments

We thank members of Kristin Scott’s laboratory, Julianne Peláez, Teruyuki Matsunaga and Hiromu Suzuki for helpful comments and suggestions; we also thank Junke Li for help with fly husbandry. This work was supported by National Institutes of Health [R01DC013280 to K.S].

## Notes

### Competing Interest Statement

The authors have declared no competing interest.

