## Supplemental Figures for "Taste adaptations associated with host-specialization in the specialist *Drosophila sechellia*"

### Supplementary Figures

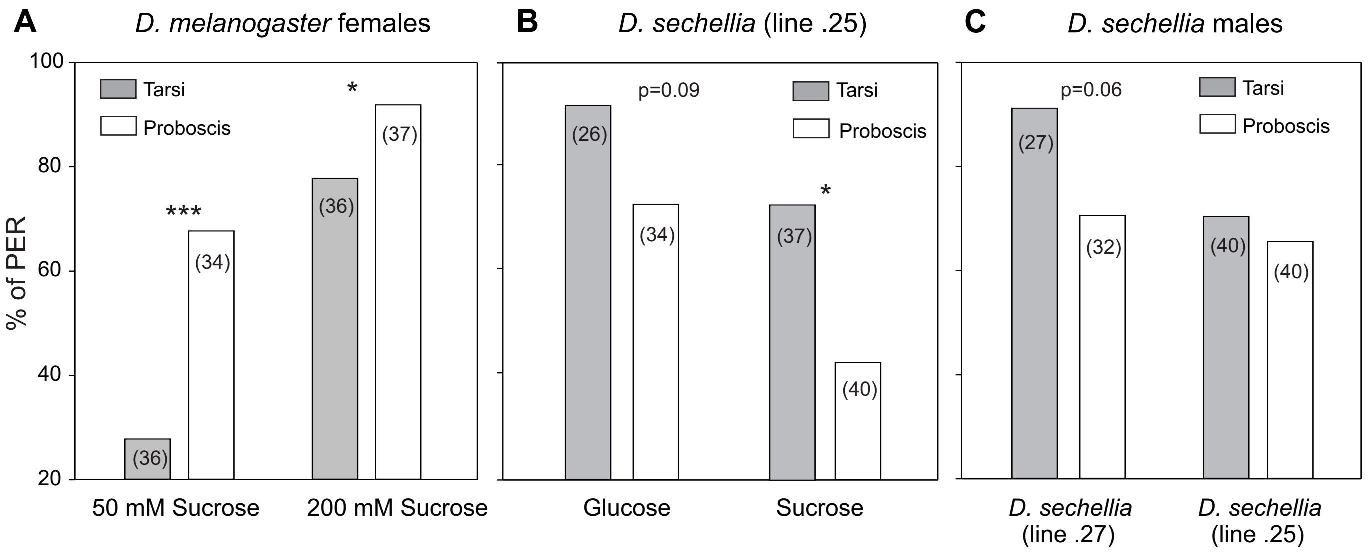


**Figure S1 (related to Fig. 1): PER of *D. melanogaster* to low sugar concentrations (A) and of different strains and sexes of *D. sechellia* (B-C).** Twenty-four hours food-deprived mated flies were stimulated three times with a drop of sugar solution applied to the tarsi or the proboscis, but were not allowed to drink, as in Figure 1. Each fly was tested with only one condition; data show the proportion of flies that extend their proboscis at least once. **A.** Proboscis stimulation evoked stronger PER than tarsi stimulation in female *D. melanogaster* at low sugar concentrations (Fisher Exact tests, ***$p<0.005$, *$p<0.05$). **B-C:** Data was obtained from a different strain of *D. sechellia* (14021-0248.25). Females also tended to show stronger PER to tarsi than to proboscis stimulation (Fisher Exact tests, *$p<0.05$ and $p=0.09$ for stimulation with 1 M sucrose or glucose, respectively). **C:** *D. sechellia* males (left: strain used throughout the manuscript, 14021-0248.27) stimulated with 1 M glucose also showed a tendency for stronger PER to tarsi stimulation (Fisher Exact tests, $p=0.061$). Also shown are results obtained from males of the strain 14021-0248.25. These results show that taste-organ PER differences are species- rather than strain-specific

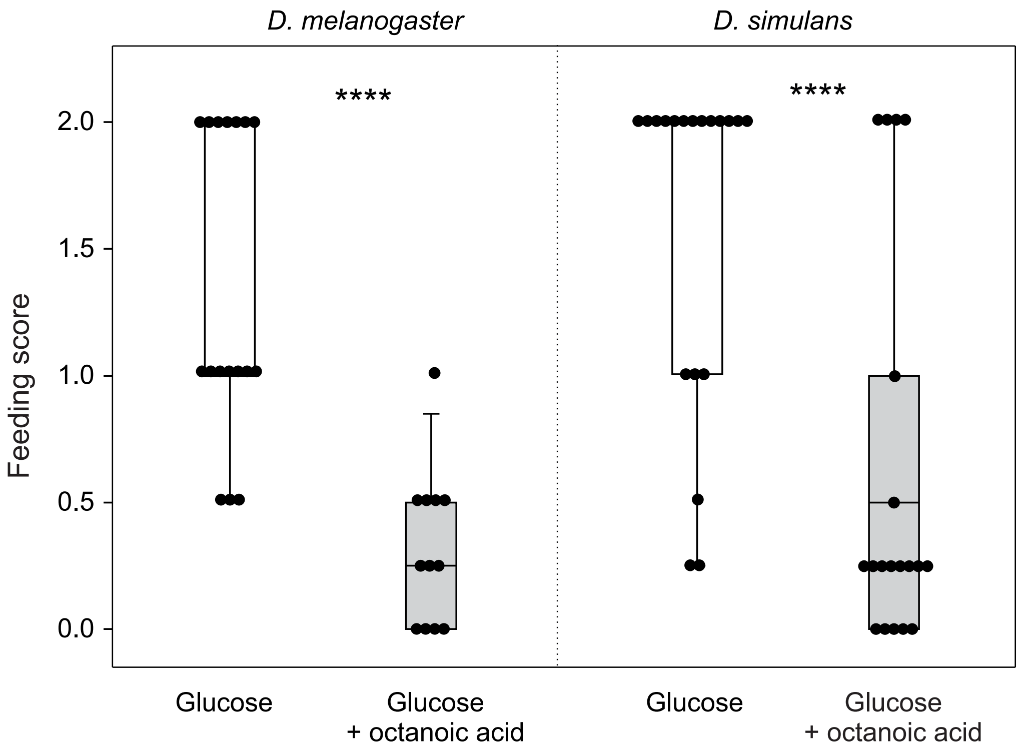


**Figure S2 (related to Figure 3): Reduced feeding on solutions containing octanoic acid is not due to lethality.** Feeding scores of individual *D. melanogaster* and *D. simulans* flies offered 750 mM glucose or glucose 750 mM + 100 mM octanoic acid. Flies were food-deprived as before and assayed singly for 30 minutes, recording viability every 5 minutes. Flies that were dead at the end of the 30-minute period (4/16 and 1/19 *D. melanogaster* and *D. simulans* offered glucose + octanoic acid) were discarded. The remaining flies were frozen and scored as before, and that single feeding score constituted an experimental unit. Symbols are individual feeding scores, boxes represent the 25% and 75% quartiles, the horizontal line inside boxes represents the median, and the whiskers indicate the 10 and 90% quartiles. Both species fed more on glucose than on glucose + octanoic acid (Mann-Whitney U tests, ****$p<0.001$), in accordance with Fig. 3.

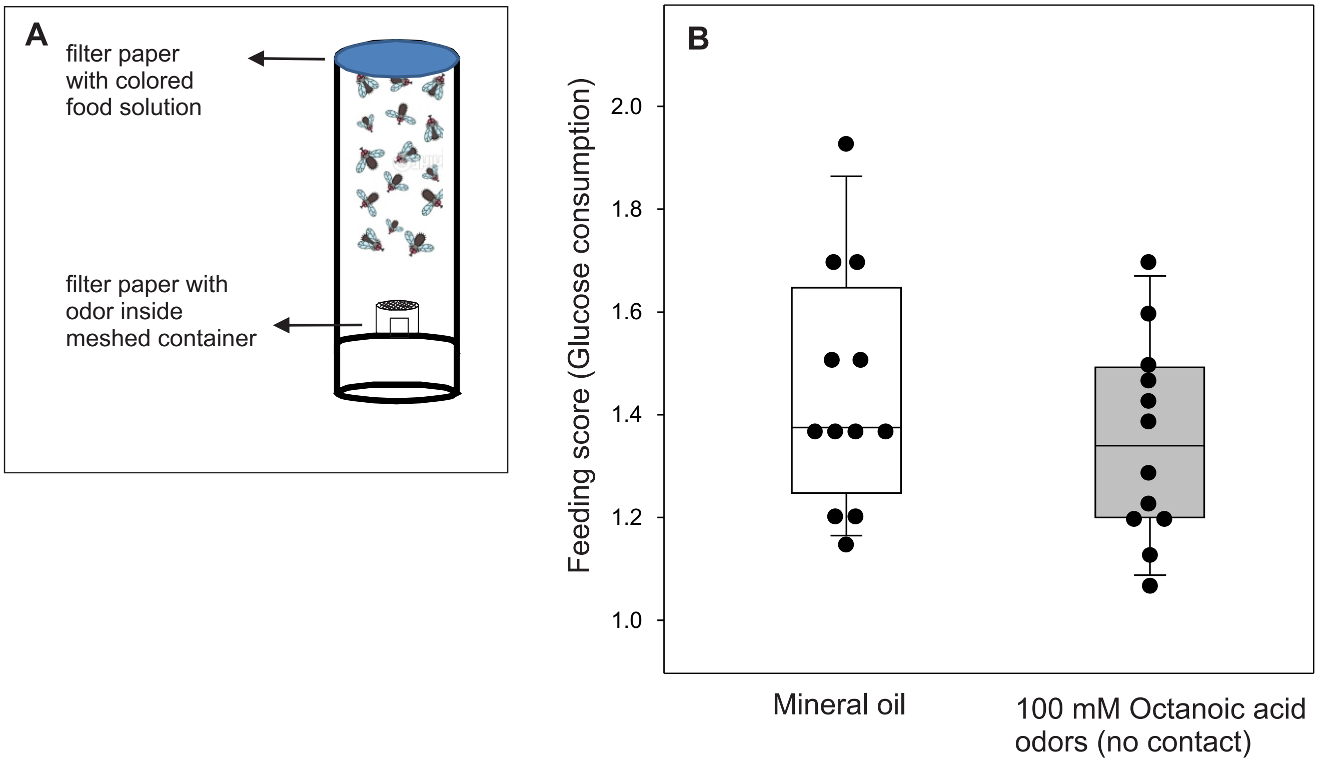


**Figure S3 (related to Fig. 3): (A)** Schematic representation of the feeding assay where flies were exposed to an odorant (10 µl on filter paper) or the solvent control (10 µl of mineral oil) with no contact (following methods described in Reisenman and Scott, 2019). **(B)** *D. melanogaster* flies consume similar amounts of 750 mM glucose in absence (white box) or presence of octanoic acid vapors (10 µl of 100 mM loaded on filter paper, gray box; Mann-Whitney U tests, $p>0.05$, $n=12$ in each group). Boxes indicate the 25% and 75% quartiles, the horizontal line inside boxes indicates the median, and the whiskers indicate the 10 and 90% quartiles; circles indicate individual datapoints. This indicates that the taste of fatty acids, not its smell, mediates feeding avoidance.

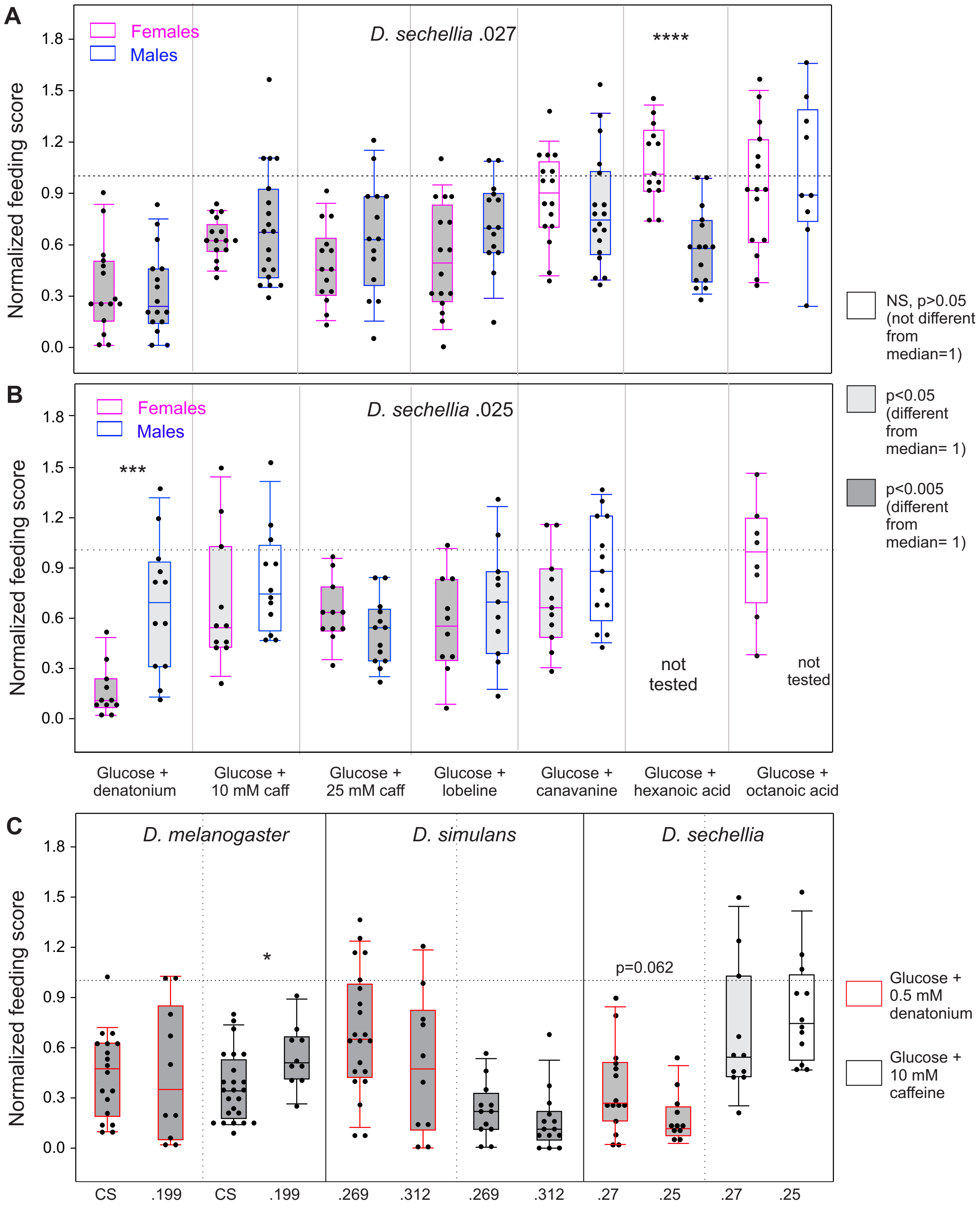


**Figure S4** (see next page)

**(from previous page)**

**Figure S4 (related to Figure 4): The reduced aversion of *D. sechellia* to bitter compounds is similar across sexes and strains. A**: strain 14021-0248.27, used throughout the manuscript, n=9-19; **B**: strain 14021.0248.25, n=8-14. Feeding scores were normalized as described in Figure 4; compound concentrations as in Fig. 4. Data represent individual data-points (symbols), median values (horizontal line within the box), the 25 and 75 percentiles (outer box lines), and the 10 and 90 percentiles (whiskers). The horizontal dotted lines at 1 indicate similar consumption on control (glucose only) and test solutions (glucose + bitter or fatty acid compound). Bar shadings indicate differences from the expected median=1 (one-sample Signed rank tests). Differences between sexes of *D. sechellia* were in most cases not statistically different (Mann-Whitney U tests, $p>0.05$). In **A**, females offered 750 mM glucose + 100 mM hexanoic acid fed significantly more than males (asterisks, $p<0.001$); males tested with 750 mM glucose + 50 mM (0.8% vol/vol) hexanoic acid still consumed less than females tested with 750 mM glucose + 100 mM hexanoic acid (normalized median=0.76, n=6, not shown). Males showed significantly reduced responses to 750 mM glucose + 100 mM hexanoic acid ($p<0.005$) but not to glucose + 100 mM octanoic acid ($p>0.05$, one-sample Signed rank tests). In **B** not all compounds could be tested; aversion to denatonium was stronger in females (asterisks). Both strains of *D. sechellia* had consistent significant aversion to denatonium, 25 mM caffeine, and lobeline; similarly, both strains lost the aversion to fatty acids (and for the most part to canavanine). **C:** Responses from two different strains of *D. melanogaster*, *D. simulans* and *D. sechellia* to 750 mM glucose with 0.5 mM denatonium (red outline boxes) or 10 mM caffeine (black outlined boxes) added (data for CS, .269 and .27 strains from Figure 4). Additional strains (all obtained from the *Drosophila* species stock center) used were: *D. melanogaster* 14021-0231.199 and *D. simulans* 14021-0251.312 (n=10-17/strain). For each species, the responses to solutions containing denatonium were not different between strains (Mann-Whitney U tests, $p>0.05$); the responses of the two *D. melanogaster* strains to caffeine were different in intensity (Mann-Whitney U test, $p<0.05$), but both strains showed aversion (one sample signed rank tests, $p<0.005$, dark gray shade). The responses of *D. sechellia* to caffeine were slightly aversive in one of the strains (one sample signed rank tests, $p<0.05$, light gray bar; white bar: $p>0.05$); responses between strains were not different from each other (Mann-Whitney U test, $p>0.05$).

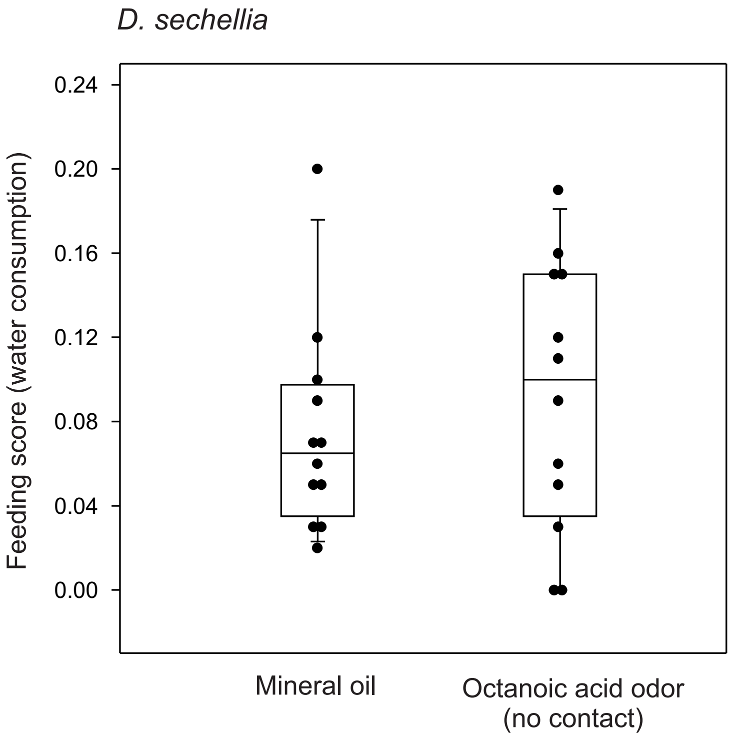


**Figure S5 (related to Figure 6): Octanoic acid odors do not increase water consumption in *D. sechellia*.** *D. sechellia* flies were offered water dyed blue in presence of octanoic acid vapors (10 µl of 100 mM on filter paper) or the solvent control (10 µl of mineral oil on filter paper) during 30 minutes (see Fig. S3A), and scored as before. Flies consumed similar amounts of water in presence or absence of octanoic acid odors (Mann-Whitney U test, $p>0.05$, $n=12$ in each group).
